# Astrocytes and neurons encode natural stimuli with partially shared but distinct composite receptive fields

**DOI:** 10.1101/2025.10.06.680791

**Authors:** Sihao Lu, Simon R. Schultz, Andriy S. Kozlov

## Abstract

Astrocytes are increasingly recognized as active participants in sensory processing, but whether they show selective responses to stimulus features, analogous to neuronal receptive fields, is not yet established. To address this, we used two-photon calcium imaging in the auditory cortex of anesthetized mice during presentation of natural ultrasonic vocalizations. Our aim was to compare astrocytic responses with those of neighboring neurons and to determine whether astrocytes exhibit feature-selective receptive fields. Event detection showed that astrocytic calcium activity is highly heterogeneous, but only a minority of events were consistently stimulus-linked. To examine this stimulus-driven subset, we estimated receptive field features using maximum noise entropy modeling and compared them with those of concurrently recorded neurons. Despite qualitative similarities in receptive-field features, analysis of modulation spectra and principal angles showed that astrocytic and neuronal receptive fields overlap but occupy distinct regions of feature space. This indicates that astrocytes and neurons encode partially shared, but not identical, dimensions of the sensory stimulus. Our findings indicate that astrocytes encode diverse sensory features, providing an additional contribution to neuronal encoding. This suggests that astrocytic calcium activity is not simply a reflection of neuronal firing, but instead represents a distinct component of cortical sensory processing.

**New and noteworthy:** We used two-photon imaging to record calcium activity in astrocytes and neighboring neurons during presentation of natural ultrasonic vocalizations. We show that astrocyte activity is highly heterogeneous across spatial and temporal scales. Further analyses indicate that a subset of astrocyte calcium activity is stimulus-linked and encodes dimensions of the stimulus that partially overlap but are not identical to those encoded by neurons.

## 1 Introduction

The study of sensory systems has largely eschewed astrocytes in favor of neurons. This is in large part due to the historical lack of tools to study them. Electrophysiological recording proved to be unexciting when applied to these cells as they do not exhibit the characteristic spikes seen in neurons. On the other hand, astrocytes display excitability in the form of an increase in calcium concentration [1], which has been studied extensively in the context of neural circuit modulation [2, 3, 4, 5, 6]. Their morphological characteristics put them in an ideal position for this role: astrocytes possess highly ramified processes that form a wide network through which they can interact with neurons [7]. The development of genetically encoded calcium indicators [8, 9] along with the advent of multi-photon imaging techniques [10, 11] has opened the door to an extensive study of these cells during normal functioning *in vivo*.

While much work has been devoted to studying astrocytes in the context of sleep [12, 13, 14, 15], memory [16, 17, 18, 19, 20], learning [21, 22, 23] and brain homeostasis [24], few studies have explored their role in sensory coding [25, 26, 27]. One argument against the importance of astrocyte calcium signaling is the fact that the timescale of the response is relatively slow compared to neuronal spiking activity. On the other hand, their ability to respond to visual stimuli in a selective manner [25] points towards a possible role in sensory processing. In recent work, astrocytes in the hippocampus have been shown to encode spatial information [28] through integration of past events [29] and through reward association [30].

Likewise, the development of computational methods to characterize sensory response properties has been largely directed toward neuronal signaling [31, 32, 33, 34]. The binary nature of neuronal spikes proved to be an appealing property which stimulated the proliferation of receptive-field estimation methods. The early methods [31, 32] were instrumental in our ability to uncover the basic coding properties of neurons throughout the brain. Crucially, these methods are based on key assumptions about the distribution of the stimuli used. The principal assumption is that the distribution is circularly symmetric. When playing simple stimuli such as pure tones or oriented gratings, the experimenter can carefully sample the space of all possible stimuli to obey this assumption. But this assumption does not hold with natural stimuli such as animal vocalizations or realistic visual scenes that exhibit a strong covariance structure. Using theoretical work in digital signal processing, methods rooted in information theory were developed to address this problem [35, 36, 37]. These methods have facilitated further discoveries in the sensory encoding of natural, ethologically relevant stimuli [38, 39, 40, 41].

Unlike the stereotypical neuronal spiking, astrocyte calcium activity is highly heterogeneous throughout a single cell, often exhibiting waves traveling across the entire cell in combination with point sources located in the fine processes of the microdomain [42]. This is not amenable to traditional region-of-interest (ROI) based analysis methods. Multiple, distinct calcium events could be occurring within the same ROI, such as a traveling wave covering a point-like event in the microdomain. It is precisely these events in the microdomains which have been implicated in sensory coding [27, 43]. Only recently have methods been developed to disentangle distinct events from astrocyte calcium imaging data [44].

In this work, our objective is to expand on the existing literature by studying the *in vivo* response properties of subcellular compartments within astrocytes in response to natural vocalizations in the mouse auditory cortex. In addition, we compare these properties with those of concurrently observed neurons. We show that astrocytes exhibit composite receptive fields, much like neurons. The features within these astrocytic receptive fields are not spectrotemporally distinct from those of neurons, and at the population level, they span overlapping but different subspaces of the full sensory space compared to neurons.

## 2 Materials and Methods

### 2.1 Ethical Statement

All procedures were carried out under the terms and conditions of licences issued by the UK Home Office under the Animals (Scientific Procedures) Act 1986. Adult, female C57BL/6 mice (Charles River, Margate, UK) were used for recordings (N=9). Mice were aged between 3-6 weeks at time of injection and 5-10 weeks during imaging experiments.

### 2.2 Auditory stimulation

Animals were stimulated with a series of natural vocalizations. Vocalizations were manually extracted from recordings obtained through an online database ^1^. Continuous segments of USVs were manually extracted using Audacity ^2^. Each segment was 8 seconds with an additional 1 second of silence inserted at both the beginning and end. These were concatenated to a 150-second stimulus, and 20 seconds of silence was added to the start to calculate baseline fluorescence pre-stimulus. Stimuli were subsequently cleaned using the Noise Reduction function and their intensity normalized to 60 dB SPL. Stimuli were played at 250 kHz through an audio interface (Avisoft UltraSoundGate Player 116) and an ultrasonic dynamic speaker (Avisoft Vifa). Playback was synchronized with image acquisition through a TTL pulse sent from the audio interface into the trigger port of the imaging system. The concatenated stimuli were played for 20 repetitions.

### 2.3 Intra-cortical AAV injection

A virus mixture consisting of either (1) 0.9 µL AAV5-gfaABC1D-cyto-GCaMP6f and 0.3 µL sterile saline, or (2) 0.3 µL AAV5-gfaABC1D-cyto-GCaMP6f, 0.3 µL AAV5-gfaABC1D-Lck-GCaMP6f and 0.3 µL AAV9-hSyn-jRGECO1a was front filled into a glass micropipette.

The injection pipette was inserted into the brain at a depth between 250 and 700 µm and left to rest for 5 minutes. Subsequently, between 200 and 1500 nL of the viral mixture was manually injected at roughly 300 nL/min using a syringe connected to the pipette via an electrode holder with a Luer-Lock attachment (World Precision Instruments, #MPH6S) mounted onto a stereotaxic frame. After injection, the pipette was allowed to rest for 5 minutes to prevent virus efflux when raising the pipette. Finally, the wound was closed using resorbable sutures (Ethicon, 6-0 Vicryl).

Throughout the procedure, the temperature of the animal was monitored and kept constant at 35°C using an electric heating pad. After the procedure, the animal was placed in a recovery chamber at 37°C. The first three days post-surgery, carprofen was added to the drinking water of the animals (190 µL of 5 mg/kg formulation in 250 mL of water).

Approximately two weeks are required for the expression of the protein encoded by the virus [9] and no imaging was performed more than 5 weeks post-injection.

### 2.4 Glass imaging window implantation

Animals were anesthetized using a mixture of fentanyl (0.05 mg/kg), midazolam (5.0 mg/kg) and medetomidine (0.5 mg/kg) injected intraperitoneally. Body temperature of the animal was maintained at 36 ^°^C. The hair covering the imaging site was shaved and the tissue was removed with surgical scissors. The periosteum was removed with forceps and the cranial surface rinsed with saline solution. The left temporal muscle was resected, revealing the temporozygomatic suture. Afterwards, a stainless steel 3D-printed headplate (Materialise nv., Leuven, Belgium) was centred over the injection location and attached using cyanoacrylate glue (UHU). The attachment was further reinforced with dental cement. A 3 mm craniotomy was performed and the dura mater was removed. Finally, a glass coverslip stack was inserted into the cranial window and secured with cyanoacrylate glue. The stack consisted of a 3-mm diameter (Warner Instruments, #CS-3R) and a 5-mm diameter circular coverslips (Agar Scientific, #AGL46R5-1) bonded together with UV-cured optical glue (Norland Products, Optical Adhesive 61).

### 2.5 Two-photon imaging

Data were collected from anesthetized, head-mounted mice using a Scientifica SliceScope equipped with a galvo-galvo scanning system and either a 20x/0.5NA or 40x/0.8NA objective (Olympus UMPLFLN20XW and LUMPLFLN40XW, respectively). The system was controlled via ScanImage software [45], and illumination was provided by a Ti-Sapphire laser (Mai Tai, Spectra-Physics) set to 980 nm. Laser dwell time was set to either 400 or 800 ns. Images of multiple astrocytes were acquired with a resolution between 0.6–1.0 µm/pixel, while single microdomains were captured at 0.2 µm/pixel. All imaging was targeted to layer 2/3 of the auditory cortex (approximately 130–250 µm from the pial surface).

### 2.6 Image processing

Image stacks were denoised with DeepInterpolation [46]. A U-Net convolutional encoder–decoder with skip connections, configured as a frame-prediction denoiser and optimized with mean-squared-error loss, was used (unet_single_1024_mean_squared_error). Training used a learning rate of 0.0001, a batch size of 5, and 64 steps per epoch. Each frame was predicted from 15 frames before and 15 frames after the target. After denoising, motion artifacts from breathing and drift were corrected with NoRMCorre available through the CaImAn package [47]. This algorithm applies piecewise rigid registration to perform motion correction of the image stack. These preprocessing steps increased signal-to-noise ratio, and supported reliable event detection and receptive field estimation.

### 2.7 Event identification

Astrocyte events were identified using the AQuA (Astrocyte Quantitative Analysis) toolbox [48, 44]. The detection threshold was determined separately for each recording (range: 2.8–6), corresponding to the minimum z-score a pixel had to exceed to be considered active. Events were required to span at least 3 pixels and last a minimum of 1 frame.

Within each imaging site, events with consistent spatial footprints and properties were grouped into Consensus Functional Units (CFUs). Events were clustered into a CFU if they shared at least 33% spatial overlap and comprised a minimum of 32 events. Co-occurrence between CFUs was defined when events occurred within two frames of each other, and CFUs were further grouped by hierarchical clustering if their co-activation was statistically significant (p < 0.001). These CFUs represent putative sources of astrocytic calcium activity. For each trial, CFU onset time was defined as the point at which the fluorescence signal reached 10% of its peak amplitude, and this was used to construct a binary activation matrix across trials.

### 2.8 Neuron segmentation

Neuronal ROIs and calcium traces were extracted using non-negative matrix factorization in the CaImAn package [49]. Parameters were set as follows: decay time = 0.45 s, autoregressive order = 1, background components = 12, merge threshold = 60%, and minimum SNR = 8. Multi-session registration was performed with the register_multisession() function, requiring components to be matched across at least 15 sessions.

### 2.9 Receptive field characterization

The spectral representation of the stimulus was computed by taking the power spectral density of the Short-Time Fourier Transform on segments of 4096 samples of the stimulus waveform with 50% overlap. The resulting spectrogram had a temporal resolution of 8.1920 ms and a frequency resolution of 61.035 Hz. After computing the spectrogram, the frequency axis was restricted to 45–110 kHz (ultrasonic vocalization range) and binned into *×* 28 linearly spaced bins. For each timepoint, we considered the 27 preceding time bins plus the current time bin, resulting in a 28 (time) × 28 (frequency) spectrogram. These 28 *×* 28 spectrograms were flattened into 784-dimensional vectors. Principal component analysis (PCA) was applied to reduce the dimensionality to 128 components, retaining 89.5% of the variance of the original stimulus.

Maximum Noise Entropy (MNE) [37] was then used to model stimulus-induced astrocyte activity. MNE models are of the form:

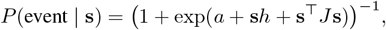

where *P* (*event* | **s**) denotes the probability of an event given a stimulus *s*, and *a, h*, and *J* are parameters optimized via gradient descent. These parameters are determined such that the predicted average activity, event-triggered average, and event-triggered covariance match those in the observed data. The receptive-field features of the model are determined as (1) the vector *h*, which represents the linear component of the response, and (2) the eigenvectors of the matrix *J*, which correspond to quadratic components of the response. These vectors are first defined in the PCA-reduced space and then projected back into the original time–frequency domain to recover the corresponding spectrotemporal features.

Stimulus-response pairs were created by averaging binary event traces of each CFU across trials and assigning the stimulus spectrogram patch immediately preceding each averaged event. The data were divided into training and test sets, with the training set comprising 70% of the total data. Models were trained using three-fold cross-validation of the training set. Prediction was performed on the remaining 30% of the data, and performance was quantified as the cosine similarity between the predicted and true responses. The significance of the observed similarity values was assessed by generating a null distribution of similarity values by randomly shuffling the stimulus-response pairs before training a new shuffled model. This randomization was performed 200 times, and each shuffled model was used to predict responses to the test stimuli. The p-value was computed as the proportion of shuffled predictions exceeding the prediction of the unshuffled model. We considered a model to be significant if its p-value was below 0.05.

### 2.10 Event clustering

We considered a subset of all the features captured by the event detection algorithm to test for the presence of distinct event classes. The features included duration, area, rise time, fall time, and circularity. This yielded a feature vector characterising each event. Properties relating to intensity of the event were omitted to avoid the effect of uneven expression of the fluorescent indicator. These feature vectors were embedded into a latent space using UMAP [50], and subsequently projected onto the first two principal components for visualization purposes. Clustering was performed using the HDBSCAN algorithm [51] in the UMAP embeddings.

### 2.11 Feature comparison

To cluster receptive field features, we first computed the modulation power spectra of each feature by applying a two-dimensional Fourier transform, yielding a joint representation of temporal and spectral modulations [52]]. These modulation spectra were then embedded into a low-dimensional latent space using the UMAP algorithm [50], a non-linear dimensionality reduction method that preserves local structure while revealing global organization. UMAP has been effectively applied to separate vocalization types across animal species [53], and to provide a compact visualization of the diversity of receptive field features [41].

### 2.12 Subspace comparison

To determine whether astrocytes and neurons are sensitive to the same dimensions within the stimulus space, we compared the subspaces encoded by their receptive field features [54]. The overlap between astrocytic and neuronal feature subspaces was quantified by computing the principal angles between the two sets. The *k*-th principal angle *θ*_*k*_ between vector subspaces 𝒰 and 𝒱 is defined as

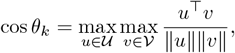

subject to the orthogonality constraint:

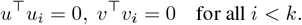

We obtained orthonormal bases for the feature sets by computing the QR decomposition, which enforces the orthogonality condition. The QR decomposition is given by

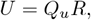

where *U* is the feature matrix, *Q*_*u*_ is the orthonormal basis spanning *U*, and *R* contains the reconstruction coefficients. Thus, *Q*_*u*_ and *Q*_*v*_ were taken as the orthonormal bases for 𝒰 and 𝒱, respectively.

For easier interpretation, the principal angles were converted from radians to degrees. A principal angle of 0^°^ indicates perfect alignment (full overlap in that dimension), whereas an angle of 90^°^ corresponds to orthogonality (no shared information in that dimension). This framework was applied both to compare astrocyte and neuronal feature subspaces, and to assess the relationship of each with the full stimulus space.

## 3 Results

### 3.1 Astrocyte calcium signals are highly heterogeneous

Calcium signaling in astrocytes shows striking heterogeneity across spatial and temporal scales; figure 1A shows an example of two events with drastically different properties within the same region. To capture this heterogeneity, we analyzed 72,615 events extracted from 396.67 minutes of calcium activity recorded using two-photon microscopy in seven imaging sites in three animals. The duration of the events spanned from a few hundreds of microseconds to the longest event, which lasted 19 seconds (Fig. 1B). Short events dominated the data, as shown by the heavily skewed distribution. Longer events were rare, but still present, as indicated by the long tail. The true distribution of event duration likely includes even shorter events, but the temporal resolution of the imaging system places a hard limit on the shortest detectable event, further restricted by the signal-to-noise ratio for shorter events. The distribution of the spatial extent of the extracted events is similarly skewed, with events smaller than 52 µm^2^ constituting half of all events and the largest event spanning nearly 5000 µm^2^ (Fig. 1C). Again, we are constrained by the spatial resolution of the microscope, which prevents us from detecting any smaller events. Moreover, the size of the field of view restricts the largest possible event that is detectable. The substantial genetic heterogeneity between astrocytes [55] led us to ask whether calcium signaling events cluster into distinct classes. To test this hypothesis, we quantified each event by its temporal and spatial properties and then applied dimensionality reduction to embed these features into two dimensions (Fig. 1C). The resulting projection revealed discrete clusters of events, implying the existence of distinct event classes.

**Figure 1:**
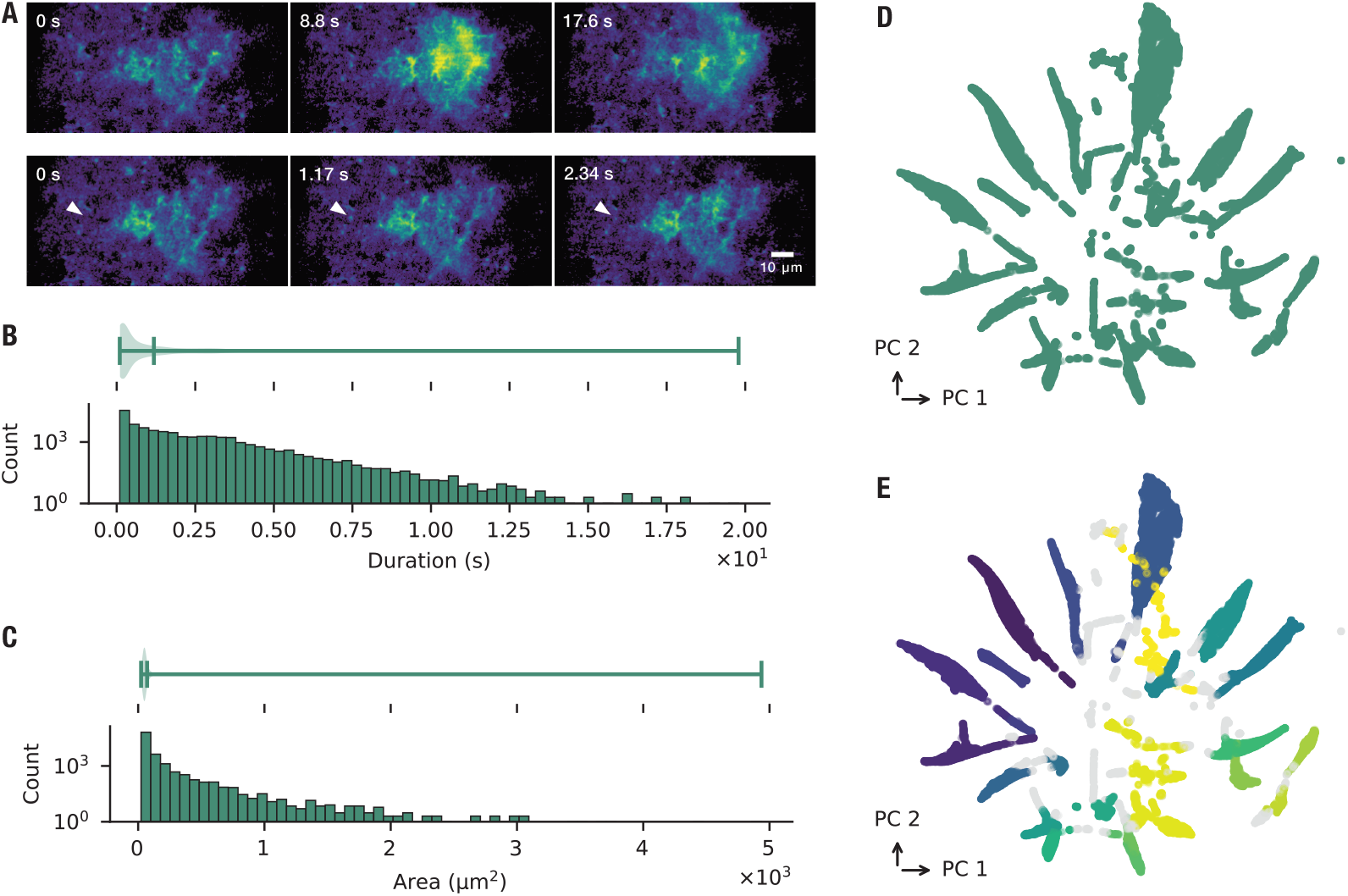
Spatial and temporal diversity of astrocytic calcium events. (**A**) Example of astrocyte events on different spatial and temporal scales. (*Top*) Onset and spread of large-scale calcium wave spread over three astrocytes, occurring in a time span of more than 17 s. (*Bottom*) The same imaging site as above. A small event can be seen (indicated by the arrow). Zero seconds in both rows signifies the start of the event. (**B**) Distribution of event duration for all of the detected events. The histogram shows counts on a logarithmic scale, and the violin plot above summarizes the overall distribution. (**C**) Distribution of area covered by event, for all of the detected events. The histogram shows counts on a logarithmic scale, and the violin plot above summarizes the overall distribution. (**D**) Dimensionality reduction of five event properties (duration, area, rise time, fall time, and circularity) of all events. (**E**) Hierarchical clustering of projection in (D). Each colour indicates a separate cluster, and gray indicates events which do not belong to any cluster. *n* = 72,615 events. Total duration of recordings: 396.67 minutes.

These clusters were quantitatively validated by applying the HDBSCAN algorithm which identified a total of 24 clusters in the data (Fig. 1D).

The heterogeneity of calcium dynamics extends to events within a single astrocyte. From high-resolution microdomain imaging, we extracted 56,225 events recorded from five imaging sites in three mice (total recording time: 255 minutes). These events exhibited a broad range of durations and spatial footprints, closely matching the heterogeneity observed at the whole-cell level (Fig. 2A, B). To further examine this variability, we quantified event properties including area, circularity, rise time, fall time, and duration, and projected them into a low-dimensional space using dimensionality reduction (Fig. 2C). Even within restricted subcellular regions, calcium events were highly diverse and distributed across a continuum of properties, without evidence for discrete clusters.

**Figure 2:**
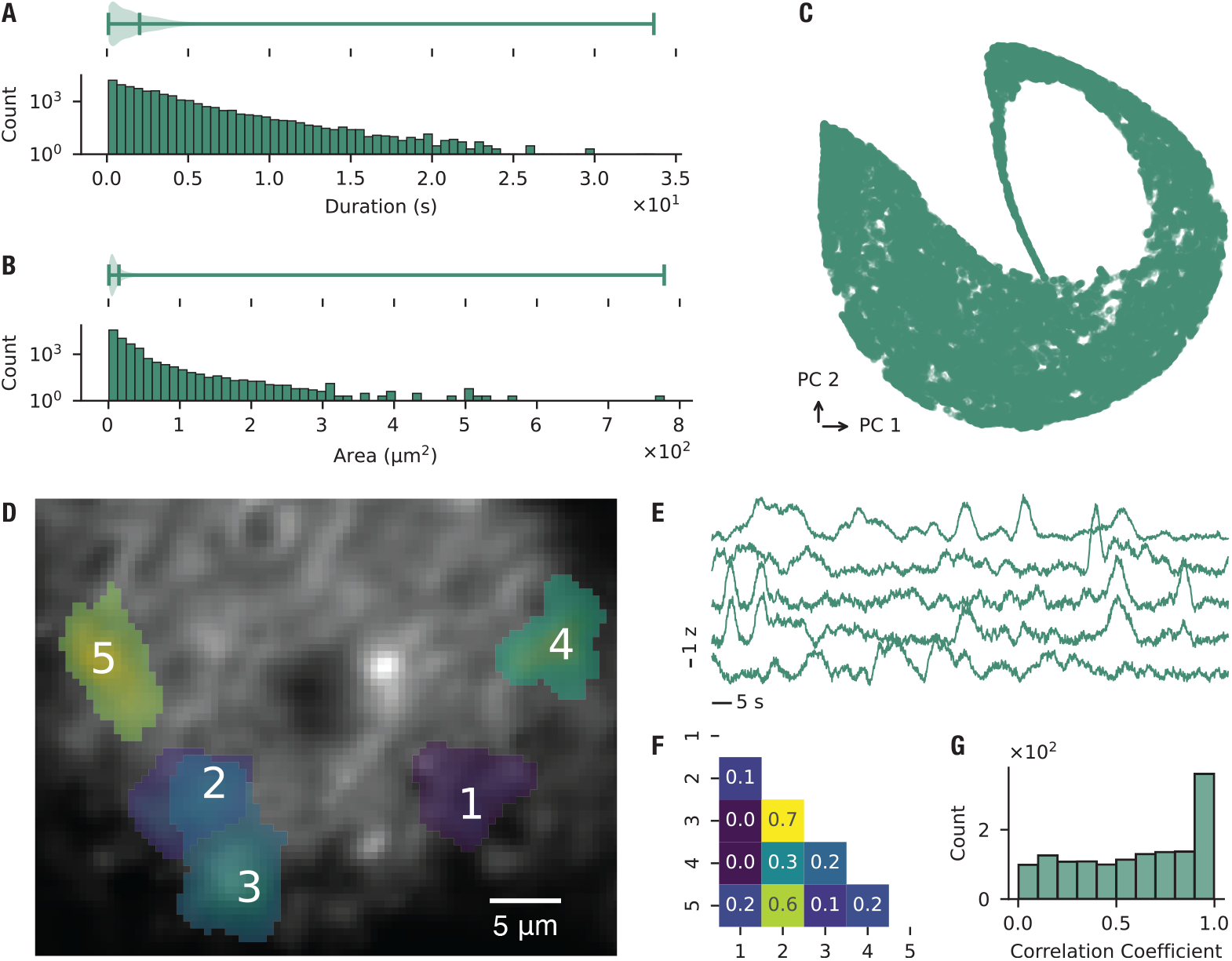
Microdomain calcium events exhibit diverse properties and weak correlations. (**A**) Distribution of event duration for all of the detected events. The histogram shows counts on a logarithmic scale, and the violin plot above summarizes the overall distribution. (**B**) Distribution of area covered by event, for all of the detected events. The histogram shows counts on a logarithmic scale, and the violin plot above summarizes the overall distribution. (**C**) Dimensionality reduction of five event properties (area, circularity, fall time, rise time, and duration) of all events. (**D**) Example of microdomain imaging. Highlighted patches denote Consensus Functional Units (CFUs), defined as loci of consistent activity across repeated stimulus presentations. (**E**) Example mean traces of events denoted in (D). (**F**) Correlation between the traces shown in (E). (**G**) Distribution of all pairwise correlations of detected event areas across all microdomain recordings. *n* = 56,225 events, 22 CFUs, 5 imaging sites, 3 mice. Total duration of recordings: 255 minutes.

Because clustering did not capture clear classes of microdomain events, we next asked whether stimulus-linked patterns of activity could be detected across trials. Within individual imaging sites, we identified loci of consistent activity across repeated stimulus presentations, referred to as Consensus Functional Units (CFUs; Fig. 2D). Example traces from several CFUs within a single field of view illustrate the diversity of temporal profiles (Fig. 2E). We computed pairwise correlations between CFUs within each microdomain and then aggregated across all recordings; this analysis confirmed that stimulus-evoked responses were generally weakly correlated or independent (Fig. 2F, G). Together, these results show that astrocytic microdomains generate a rich and heterogeneous repertoire of calcium signals.

### 3.2 Astrocytes have composite receptive fields

To characterize astrocyte calcium responses in the auditory cortex, we presented USVs to nine anesthetized mice and extracted calcium events from a total of 24 imaging sites (Fig. 3A). Stimulus-specific events were identified by grouping events that displayed a consistent spatial footprint and event properties across 20 repeated presentations of the stimulus. These loci are referred to as Consensus Functional Units (CFUs), following the terminology of the AQuA framework. Across all recordings, we detected 231 CFUs. Within a given imaging site, however, only a small fraction of recorded events were assigned to CFUs, indicating that most activity was not stimulus-driven. The auditory stimulus and its PCA reconstruction are shown in Fig. 3B, and representative CFU activation patterns across trials in Fig. 3C. Having defined CFUs as stimulus-linked loci of activity, we next estimated their receptive fields using the maximum noise entropy (MNE) method, previously applied to neuronal responses [38, 40, 41]. Model performance was evaluated on held-out data by calculating cosine similarity between predicted and recorded responses. A total of 59 CFUs passed our significance threshold and were considered for further analysis, yielding a mean cosine similarity of 0.63 ± 0.1 (Fig. 3D). Across these CFUs, the model identified 103 significant features, with an average of 2.12 ± 1.78 inhibitory and 0.03 ± 0.18 excitatory features per CFU (Fig. 3E). Examples of recovered receptive-field features are shown in Fig. 3F. Together, these results indicate that astrocytic calcium activity can be stimulus-linked and is selectively modulated by distinct features within natural vocalizations.

**Figure 3:**
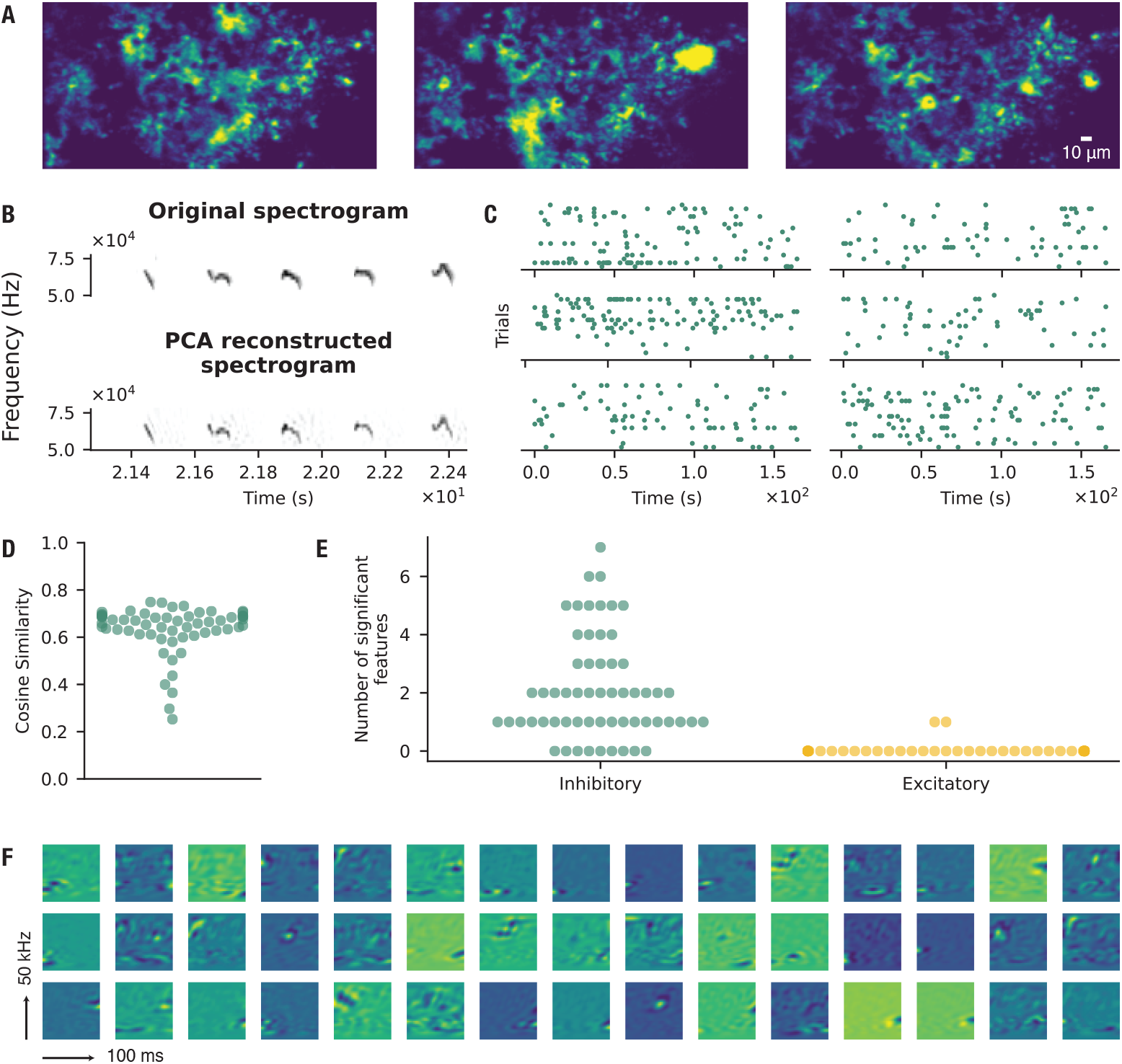
Astrocyte calcium signals in auditory cortex exhibit composite receptive fields. (**A**) Representative astrocyte calcium signaling events visualized as maximum-intensity projections over a 42.5-second recording window. (**B**) Original spectrotemporal representations of ultrasonic vocalizations (USVs) alongside their principal component analysis (PCA) reconstructions used for auditory stimulation. (**C**) Examples of binary matrices of CFU activation during auditory stimulation. Rows represent individual trials, with entries indicating event onset times. (**D**) Cosine similarity values for fitted models (*µ* = 0.63, *σ* = 0.10). Cosine similarity is computed between model predictions and held-out test data. Higher values indicate better generalization performance of the model. Each point corresponds to a model trained on one CFU. (**E**) Number of significant features identified from each model and classified as inhibitory (decreasing event probability) or excitatory (increasing event probability). *µ*_in_ = 2.12, *σ*_in_ = 1.78; *µ*_ex_ = 0.03, *σ*_ex_ = 0.18. *n* = 59 CFUs, 24 imaging sites, 9 mice. (**F**) Examples of receptive-field features estimated from astrocyte calcium activity in response to auditory stimulation.

### 3.3 Astrocytic and neuronal receptive fields occupy overlapping but distinct regions of feature space

Neurons have long been recognized to respond selectively to sensory stimulation. To compare astrocytic and neuronal receptive fields, we imaged 24 sites in the auditory cortex of 9 anesthetized mice during repeated presentation of USVs. From these recordings, we identified 59 CFUs and 317 neurons. Examples of concurrently acquired calcium signals from astrocytes and neighboring neurons are shown in Fig. 4A. Neuronal receptive fields were estimated using the maximum noise entropy (MNE) method, as applied above for astrocytes. Of the 317 neurons, 141 yielded models that generalized significantly to new data. Model performance on held-out data, quantified by cosine similarity, is shown in Fig. 4B. Across these neurons, we identified 249 significant features, with an average of 1.71 ± 1.41 inhibitory and 0.06 ± 0.23 excitatory features per neuron (Fig. 4C, D).

**Figure 4:**
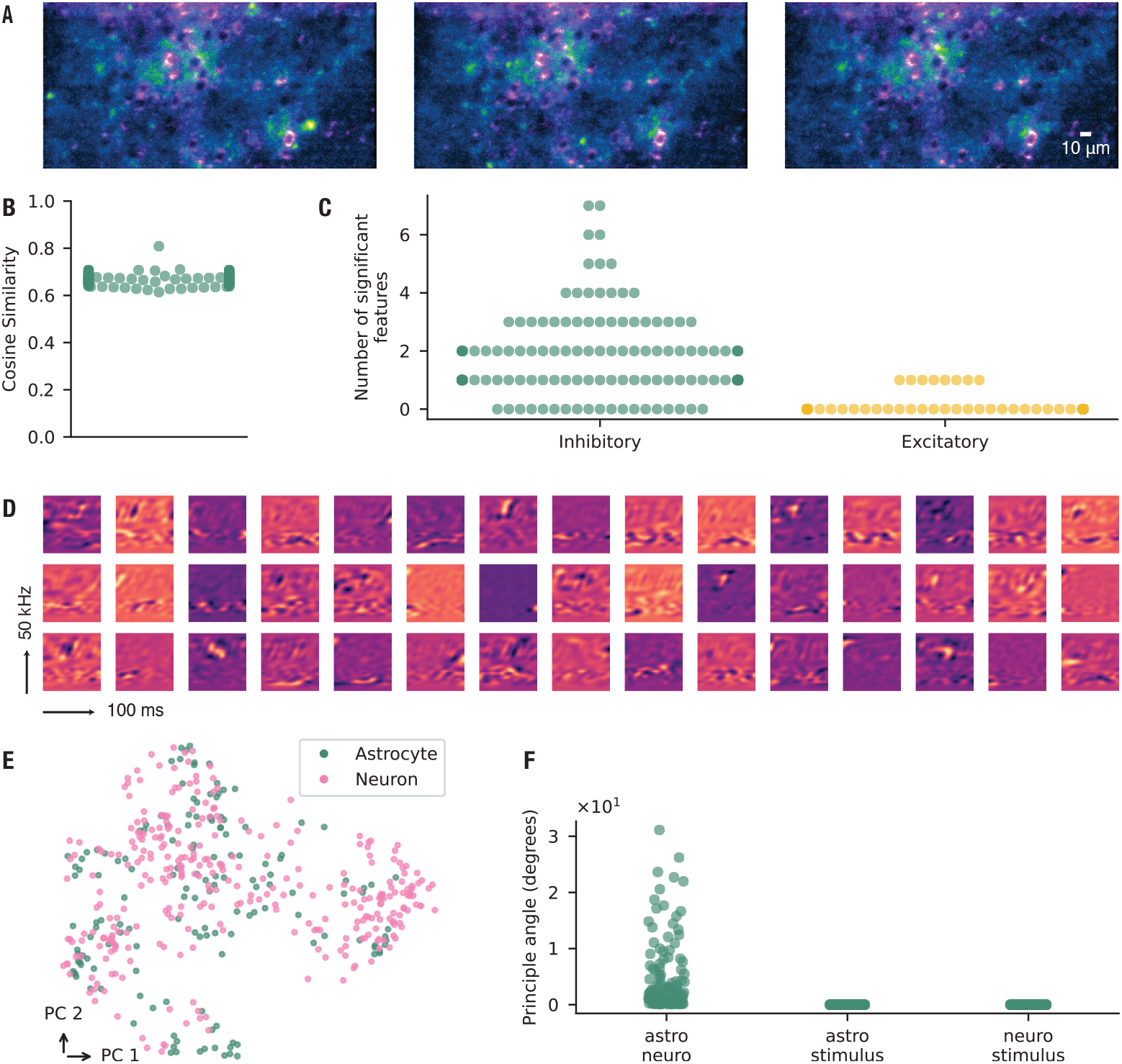
Astrocytic and neuronal receptive fields encode overlapping but distinct feature spaces. (**A**) Representative astrocyte calcium events (green) and neuronal calcium activity (magenta) shown as maximum-intensity projections over a 42.5-s recording window. (**B**) Cosine similarity values for all fitted models, calculated between predicted and recorded responses (*µ* = 0.67, *σ* = 0.02). (**C**) Number of significant features identified from the model and classified as inhibitory (decreasing event probability) or excitatory (increasing event probability). Each point corresponds to a model trained on one neuronal CFU. *µ*_in_ = 1.71, *σ*_in_ = 1.41; *µ*_ex_ = 0.06, *σ*_ex_ = 0.23. (**D**) Examples of receptive field features estimated from neuronal calcium activity in response to auditory stimulation. (**E**) UMAP projection of modulation spectra of all estimated receptive field features for astrocytes and neurons. (**F**) Principal angles between the subspace encoded by the astrocyte receptive field features and the neuron receptive field features (astro-neuro). Also shown are the principal angles between the subspace encoded by the astrocytes or neurons, and the full stimulus space respectively (astro-stimulus, and neuro-stimulus). *n* = 59 CFUs, 141 neurons, 17 imaging sites, 7 mice.

Qualitatively, the receptive-field features of astrocytes and neurons appeared similar. To quantify this relationship, we embedded their modulation spectra into a low-dimensional representation to test whether astrocytic and neuronal features formed distinct clusters. No clear separation was observed (Fig. 4E). Although astrocytic and neuronal receptive fields appeared qualitatively similar (Fig. 4E), we next asked whether they in fact encoded the same subspace. To test this, we applied principal angle analysis: if one feature set were fully contained within the other, the principal angles would approach zero. Instead, we observed a substantial number of non-zero angles, indicating that the two populations occupy overlapping but distinct regions of feature space (Fig. 4F). As a control, we also computed principal angles between each feature set and the full stimulus space, confirming that both astrocytic and neuronal receptive fields form subsets of the overall stimulus space.

## 4 Discussion

The study of astrocytes and other glia has enjoyed a rise in popularity in recent years [56, 57], aided by the advent of powerful tools such as GECIs together with steady advances in imaging technology. These tools facilitate the systematic study of glial function, but many questions remain. For example, different genetic subtypes of astrocytes may have specialized roles in the context of sensory processing [58, 59], but this has not been systematically explored. In addition to existing evidence [2, 3], new mechanisms by which astrocytes modulate neuronal circuits are being uncovered [56, 60, 61]. However, the specific roles of the various calcium sources remain unclear [62]. Another unresolved question is whether information is represented primarily at the population level across many astrocytes or within the microdomains of individual cells.

In the present work, we show that astrocyte calcium activity in the mouse auditory cortex is highly heterogeneous, and that a subset of this activity is modulated by multiple features within naturalistic stimuli. These data parallel those reported in neurons [41]. Crucially, we show that the receptive-field features identified in astrocytes and neurons occupy overlapping yet distinct regions of feature space. Thus, some stimulus dimensions are represented preferentially by astrocytes, while others are only encoded by neurons. These results argue against the idea that astrocytic calcium activity simply reflects a direct, linear consequence of neuronal firing. Rather, astrocytes have been shown to exhibit integrative properties that allow them to modulate neuronal circuits [63, 64, 29]. To further dissect these mechanisms, it will be important to measure directly how neuronal activity drives astrocytic responses at scale, and to determine how distinct signaling pathways such as sodium signaling [65], potassium buffering [66, 67], norepinephrine uptake [68, 69, 70], and other modulatory inputs contribute to shaping astrocytic function. We propose that to support their functions in circuit modulation, astrocytes must be capable of detecting and responding selectively to specific stimulus features. Therefore, a more detailed analysis of the functional relevance of the different event classes to sensory processing will be required.

Our study is limited by the imaging tools used: subcellular calcium signals may be difficult to detect, and very brief events may not be captured at the frame rates employed or due to the kinetics of the GECI. The observed distributions of event durations and sizes suggest that we have not yet reached the limits of what astrocytes can generate. Furthermore, our experiments were performed under anesthesia, which is known to alter the physiology of both neurons and astrocytes [71]. Astrocytic calcium activity in awake animals can be faster, more spatially extensive, and strongly modulated by the behavioral state [12], suggesting that the repertoire of responses described here represents only a portion of their full physiological capacity. Future studies that combine higher-speed or more sensitive indicators with large-scale recordings of both neurons and astrocytes in awake animals will be needed to define the principles by which glial and neuronal activity interact to shape cortical representations.

## Acknowledgments

The authors thank Dr. Amanda Foust for help with microscope troubleshooting, Dr. Mary Ann Go for guidance on surgical techniques, and Dr. Yu Liu for assistance with viral injections.

We thank Baljit Khakh for providing the constructs pZac2.1 gfaABC1D-lck-GCaMP6f (Addgene viral prep #52924-AAV5; RRID:Addgene_52924) and pZac2.1 gfaABC1D-cyto-GCaMP6f (Addgene viral prep #52925-AAV5; RRID:Addgene_52925). We also thank Douglas Kim and the GENIE Project for providing pAAV.Syn.NES-jRGECO1a.WPRE.SV40 (Addgene viral prep #100854-AAV9; RRID:Addgene_100854).

Present address for Sihao Lu: Laboratory of Sensory Neuroscience, The Rockefeller University, New York, NY, USA; Howard Hughes Medical Institute, The Rockefeller University, New York, NY, USA.

## Author contributions

S.L and A.S.K conceived and designed research. S.L. analyzed data. S.L performed experiments. S.L and A.S.K interpreted results of experiments. S.L prepared figures. S.L drafted manuscript. S.L, S.R.S, and A.S.K edited and revised the manuscript. S.L, S.R.S, and A.S.K approved the final version of the manuscript.

## Data availability

Raw imaging data in addition to denoised image stacks are available upon reasonable request. The code used for the analyses in the manuscript can be found on GitHub: https://github.com/Sihao/astro-neuro-rf.

## Footnotes

1 https://mousetube.pasteur.fr

2 http://audacityteam.org

